# A comprehensive investigation of m^6^A regulators prognostic value and associated molecular pathways in breast invasive cancer

**DOI:** 10.1101/2021.06.07.447359

**Authors:** Turan Demircan, Mervenur Yavuz, Sıddıka Akgül

## Abstract

Breast invasive cancer (BIC) is one of the most commonly observed and the deadliest cancer among women. Despite the progress that has been made in improving breast cancer outcomes by the development of advanced treatment options, due to the heterogeneity and complexity of the disease, more studies are required to explore underlying molecular mechanisms of breast cancer which may provide useful insights to overcome the constraints related to current therapeutic options. The goal of this study was to reveal the crucial roles of m6A regulatory proteins in BIC development using various publicly available datasets and databases. We first conducted a comprehensive analysis to depict the mutation frequency and types for m6A regulatory genes in sub-types of BIC for the evaluation of the genetic alterations landscape of breast cancer. Changes in expression levels of m6A regulatory factors were identified as the key genetic alteration in BIC. Implementation of Kaplan-Meier tool to assess the predictive value of m^6^A pathway components in BIC validated the use of VIRMA, METTL14, RBM15B, EIF3B, YTHDF1, and YTHDF3 as prognostic biomarkers of breast cancer. We then examined the enriched gene ontology (GO) terms and KEGG pathways for the tumor samples with genetic alterations in the m^6^A pathway. Dysregulation of m^6^A regulatory factors in BIC was associated with cell division and survival-related pathways such as ‘nuclear division’ and ‘chromosome segregation’ via the upregulation of the genes functioning in these biological processes and the gained overactivity of these pathways may account for poor prognosis of the disease. The performed analyses highlighted m^6^A pathway genes as potential regulators of BIC growth and as a valuable set to be utilized as clinical biomarkers in BIC disease.

## 1. Introduction

Breast cancer is the second commonly diagnosed cancer with the highest mortality rate among women all around the world [1,2] The subtypes of malignant breast invasive cancers reflect the type of the mutations and primary tumor site [3]. Breast invasive ductal carcinoma (BIDC) is the most frequently identified breast cancer and is primarily characterized by tumor formation in milk ducts [3,4]. The second most common breast cancer, breast invasive lobular carcinoma, arises from the milk-producing lobules [5]. Breast mucinous carcinoma originating from mucus-producing cancer cells is a rare type of breast cancer and classified into two subtypes. In breast invasive pure mucinous, carcinoma transformed cells are surrounded by extracellular mucin layer. On the other hand, breast invasive mixed mucinous carcinoma (BIMC) consists of cells from other types of breast cancers such as infiltrating ductal carcinoma, aside from the mucinous cells [6]. Furthermore, breast invasive cancers which cannot be classified into defined cancers due to lacking specific features are termed as not otherwise specified (NOS) [7,8]. Poor prognosis and aggressiveness of the BIC limit the benefit of current surgery, chemo and radiotherapy treatment options and therefore new therapeutic approaches are necessary to extend the toolbox for the cure of BIC by the discovery of new biomolecules to target. In this respect, reexamination of essential pathways is required to uncover their association with BIC progression and prognosis.

RNA modification refers to the chemical modification of RNA molecules co-transcriptionally or at the post-transcription level by the regulatory proteins highly conserved among various species [9]. Up to date, more than 170 modifications such as N^6^-metyhladenosine, N^1^-metyhladenosine, 5-methylcytosine, 5-hydroxymethylcytosine, pseudouridine, and inosine have been identified and these modifications have diverse roles on stability, localization, secondary structure, and decay of modified RNAs depending on the type and site of the modifications [10–13]. Although the vast majority of the characterized modifications are associated with tRNA and rRNA molecules, studies to dissect the roles of mRNA modifications particularly methylation of mRNAs have been gained momentum considering the emerging roles of these modifications on mRNA stability and translation which open the era of epitranscriptome.

mRNA methylations are controlled by dynamic reversible mechanisms and dysregulation of this mechanism is related to a broad range of diseases due to the alterations in the numerous biological processes[14–17]. Methyl group is added by a special group of proteins called “writers”, interpreted by “readers” and removed by “erasers”. Writer proteins catalyze methylation of target mRNA molecules in a context-dependent manner which can be reversed by the activity of eraser proteins through the demethylation reaction. Reader proteins which determine the fate of the modified mRNAs depending on the reader protein type and mRNA interactions, recognize methylated bases and bind to the methylated mRNAs [18].

Methylation of the adenosine base on the 6^th^ nitrogen position is termed as N^6^-methyladenosine (m^6^A) modification which is the most abundant messenger-RNA (mRNA) modification in mammals [19,20]. The distribution of m^6^A on an mRNA is not random, and in general, methylated bases are located within clusters in the 3’UTR regions and near the stop codon of the transcripts [21,22]. m^6^A modification has a substantial impact on mRNA metabolism, transcription, splicing, export, translation, degradation, stability, and localization [11,13]. Consequently, a wide range of vital biological activities at the cell and organism levels such as cell cycle, apoptosis, self-renewal, RNA metabolism, and development are affected and modulated by m^6^A modification [14,23–25]. At the molecular level, m^6^A modification destabilizes the intrinsic secondary structure, and by exposing the concealed protein-binding sites to the reader proteins, alternative splicing can be affected directly or indirectly by binding of reader proteins to methylated mRNA. Based on the evidence, reader proteins are also involved in the nuclear export of modified mRNAs to the cytoplasm by interacting with exporting complexes [13]. As another imperative role, reader proteins take part in the enhanced translation of modified mRNA molecules by binding to translation elongation factors and ribosome’s subunits [13,26]. The link between m^6^A modification and mRNA stability and degradation is established by reader proteins by recruiting mRNAs to degradation complexes or protecting methylated mRNAs from decay [13,27].

As another type of adenosine methylation, 1^st^ nitrogen of adenosine base can be methylated as well. This modification, N^1^-methyladenosine (m^1^A), occurs mostly in tRNA molecules and regulates its secondary structure. Along with tRNA, m^1^A modification is detected in mRNA molecules with the potential function of stabilizing the mRNA-protein interactions. Although the exact function of m^1^A modification in mRNA remains largely unclear, recent studies suggest that m^1^A modification may decrease the rate of translation [28]. Modification at the 5^th^ carbon of cytosine has a remarkable the methylated mRNAs’ impact on nuclear export and protein translation rate. m^5^C methylation has dual roles to regulate the translation rate in a way that methylation in 5’UTR is a signal for inhibition of translation and 3’UTR region methylation enhances the translational activity [29].

Alterations in epitranscriptome pathways have been associated with many different cancer types [30,31]. Changes in stability and translation rate of modified mRNAs fluctuate the final amount of produced proteins and thereby may account for modulation of molecular mechanisms underlying the cancer progression. Across various cancer types, binary roles of the m^6^A pathway in tumor progression or tumor suppression have been demonstrated. In glioblastoma multiforme (GBM), class IV neoplasia of glial cells, m^6^A modification leads to tumor growth, and poor prognosis of the disease by enhancing the renewal capacity of glioblastoma stem cells [32]. Elevated levels of writer proteins in the m^6^A pathway have been also correlated with increased self-renewal capacity of leukemia stem cells in acute myeloid leukemia (AML) [33] due to the increased translation rate of oncogenic and anti-apoptotic factors, such as c-Myc and PTEN, and BCL-2, respectively [34]. The oncogenic role of FTO, an eraser protein component of the m^6^A pathway, in AML is evident by targeting the tumor suppressor proteins such as ASB2 and RARA which contribute to hematopoietic cells transformation [35]. In hepatocellular carcinoma (HCC), depletion of SOCS2 mRNA by the activity of reader protein YTHDF2 and upregulated writer protein METTL3 promotes tumor formation [36]. Deficit METTL14 in HCC patients was linked to the higher metastatic capacity of cancer [37]. YTHDF2 is found to be upregulated in prostate cancer, and as shown in a recent study, knockdown of YTHDF2 inhibits cancer cell proliferation and migration [38]. Another study revealed that YTHDF2 is upregulated also in pancreatic cancer cells and increased YTHDF2 levels induce tumor progression *via* activating AKT, GSK-3B, and cylinD1. Controversially, increased levels of YTHDF2 are also a limiting factor for metastasis of pancreatic cancer cells [39].

Reduced levels of demethylating factor ALKBH5 in pancreatic cancer cells alter the migration and invasion capacity. Methylation of KCNK14-AS1, a long non-coding RNA responsible for inhibition of cell migration and invasion, makes it unstable and stimulates its degradation [40]. In a subtype of cervical cancer, cervical squamous cell carcinoma (CSCC), a highly expressed FTO protein catalyzes the demethylation of ß-catenin mRNAs to increase its expression, and as a consequence, CSCC patients become more resistant to chemotherapy and radiotherapy [41]. Diminished PHLPP2 and elevated m-TORC2 translation rate in endometrial cancer were due to these mRNAs ’decreased methylation levels. These genes are negative and positive AKT pathway regulators respectively, and induced activation of AKT pathway by dysregulation m^6^A modification is described as one of the key contributors of tumorigenesis in endometrial cancer [42].

Abnormal alterations in the m^6^A pathway have been described in breast cancer as well. Hypoxic tumor microenvironment induces ALKBH5 expression and increased levels of ALKBH5 lowers NANOG mRNA methylation to enhance NANOG mRNA stability. NANOG is an essential transcription factor in maintaining pluripotency in cancer stem cells and upregulated ALKBH5 levels increase the metastatic and invasive capacity of breast tumors through the stabilization of NANOG [43]. Wang et. al. reported that METTL3 level is raised in breast cancer which favors the anti-apoptotic activity of the cancer cells by methylation of BCL-2. [44]. Another study exploiting the TCGA datasets reported that high expression levels of methylation factor VIRMA in breast cancer patients’ samples compared to normal tissues was detected. The finding of decreased overall survival rate of patients with high expression of VIRMA in this study was notable. *In vitro* and *in vivo* assays confirm the association of VIRMA with breast cancer progression and metastasis by acting on CDK1, an important cell cycle regulatory protein [45]. As shown in another recent study, upregulated FTO in breast cancer cell lines stimulates the demethylation of pro-apoptotic BNIP3 mRNA and induces its degradation. Therefore, increased FTO level was correlated with lower survival rate, accelerated cell proliferation, colony formation, and metastasis [46].

Here, we performed a comprehensive bioinformatics analysis to explore the putative link between m^6^A, m^1^A, and m^5^C mRNA modification pathways and breast invasive cancer by utilizing several publicly available databases and BIC datasets. Molecular alterations in and expression level differences at both mRNA and protein levels of regulatory proteins belonging to these pathways in BIC were investigated. The followed in-silico experimental study revealed critical roles of epitranscriptomic pathways in BIC manifested in survival, apoptotic and proliferation-related pathways, which may open a new avenue for BIC research.

## 2. Materials and Methods

### 2.1 Used TCGA Datasets From cBioPortal

In this study, publicly available datasets on the TCGA database (http://cancergenome.nih.gov) for Breast Invasive Carcinoma (TCGA, Pan Cancer Atlas) which contains 1084 breast cancer patient samples were utilized. To analyze the mutations, structural variant, copy-number alterations, and mRNA expression z-Scores (RNA Seq V2 RSEM) for each gene, acquired data from the TCGA datasets came from the cBioportal database (https://www.cbioportal.org/) were used. Oncoprint and cancer type summary (n= 780 for Breast Invasive Ductal Carcinoma, n= 201 for Breast Invasive Lobular Carcinoma, n= 77 for Breast Invasive Carcinoma (NOS), n= 17 for Breast Invasive Mixed Mucinous Carcinoma, n= 8 for Metaplastic Breast Cancer and n=1 for Invasive Breast Carcinoma) of each gene were depicted to portrait the type of genetic alterations (missense mutation, amplification, splice mutation, deep deletion, truncating mutation, mRNA up- or down-regulation) and alteration frequency for each gene and subtypes of cancer.

### 2.2 Volcano plot and Functional Enrichment Analysis

Differentially expressed genes (DEGs, |FC|>1.5 and p<0.01) between altered (observed mutation in epitranscriptomic pathway genes) and unaltered (no mutations in epitranscriptomic pathway genes) groups for breast cancer patients were downloaded from the cBioPortal database. DEGs were displayed on a volcano plot using the ‘enhanced volcano’ R package. Top ten DEGs for both altered and unaltered groups were highlighted on the volcano plot. To explore the Gene Ontology (GO) categories enriched by DEGs between altered and unaltered groups at biological process (BP), molecular function (MF), and cellular component (CC) levels, the R language “clusterProfiler” package [47] was used. The same package was implemented to perform Kyoto Encyclopedia of Genes and Genomes (KEGG) enrichment, and Gene Set Enrichment Analysis (GSEA) of the genes belonging to altered and unaltered groups. To calculate the statistical significance, Fisher’s exact test was utilized. Enriched GO terms and KEGG pathways with adjusted *P* < 0.05 were considered statistically significant, false discovery rate (FDR) < 0.05 was used as the cutoff, and the results were visualized by ‘enrichplot’ and ‘ggplot2’ packages [48] with dotplot, barplot, ridgeplot or cnetplot visualization methods as used elsewhere [49,50]

### 2.3 *UALCAN dataset* analysis *for the mRNA expression and promoter methylation of epitranscriptomic pathway genes*

TCGA breast invasive cancer dataset on UALCAN database (publicly available at http://ualcan.path.uab.edu) was used to analyze the mRNA expression level of m^6^A pathway genes between breast cancer patients (n=1097) and normal (n=114) samples. Gene expression level by means of transcripts per million is plotted for normal and primary tumor samples for breast invasive carcinoma. Furthermore, the same dataset was used to compare and plot the mRNA level based on individual cancer stages, and the significance of comparison is depicted in the figures. Promoter methylation levels of m^6^A pathway genes were investigated using the UALCAN database as well. Beta value to compare the levels of methylation of m^6^A genes between cancer and normal groups indicates a level of DNA methylation ranging from 0 (unmethylated) to 1 (fully methylated).

### 2.4 Kaplan-Meier Plots for expression level and mutation frequency

Kaplan-Meier survival analysis method was used to compare the survival ratio within the breast cancer patients group by using the Kaplan-Meier Plotter database (https://kmplot.com/) [51]. Overall survival graphs were generated for breast cancer patients displaying low and high m^6^A pathway gene expression, and for the patients with or without mutations in m6A pathway components. P values (as Log-rank means) were used to evaluate the statistical significance of the analysis.

### 2.5 Human Protein Atlas data

The representative immunohistochemistry (IHC) images of m^6^A pathway genes were retrieved from The Human Protein Atlas [52], a publicly available database containing normal and cancer tissues labeled with a wide range of antibodies to show the spatial distribution of proteins. This high-resolution image collection was generated by using the same antibody to stain the protein of interest in both cancer and normal tissues for a particular gene. Representative IHC images of m^6^A pathway proteins in cancer and normal tissues were downloaded from the database (http://www.proteinatlas.org/). In addition to employing specific antibodies to label the target protein, hematoxylin staining was performed to provide the contrast.

### 2.6 Protein-protein interactions networks

Protein-protein interaction (PPI) networks were analyzed and visualized using the STRINGdb v11.0 (Search Tool for the Retrieval of Interacting Genes/Proteins) [53]. Genes enriched in the top significant BPs (’Nuclear division’ for altered groups and enriched in ‘Regulation of membrane potential’ for unaltered groups) were selected to generate an interaction network with m^6^A pathway genes.

### 2.7 The communality of DEGs and Enriched Terms

In order to find the unique and shared enriched GO terms among epitranscriptomic pathways for both altered and unaltered groups, jvenn, a web-based tool [54], was implemented to construct the venn diagrams. The same tool was employed to identify the commonly found and unique genes in altered and unaltered groups among m^5^C, m^6^A, and m^1^A pathways. The common 42 enriched GO terms for altered and 31 enriched terms for unaltered groups were clustered based on euclidian distance measure and complete linkage method. Plotting of common GO terms was performed using the ComplexHeat map R package) [55].

### 2.8 Statistical Analysis

R language (4.3.2) was used for statistical analysis and graphing. For pairwise comparisons between cancer and control samples, student’s t-test was performed and ANOVA test was applied to interrogate the significance among the cancer subtypes. Statistical significance was set as p < 0.05 for all.

## 3. Results

### 3.1. Mutation analyses overview of m^6^A pathway genes in breast invasive cancer

To explore the m^6^A pathway genetic alterations in BIC, comprehensive bioinformatics analysis has been performed using the R language and publicly available databases listed in the methods section. As the alterations of m^6^A pathway components at mRNA expression and DNA levels have been linked to several cancers by many recently published reports, we interrogated the genomic and transcriptomic landscape of m^6^A genes in the TCGA pan-cancer BIC dataset involving 1084 patients. We first analyzed the frequency and type of observed mutations (Figure 1) to display the alterations in m^6^A RNA modification enzymes.

**Figure 1.**
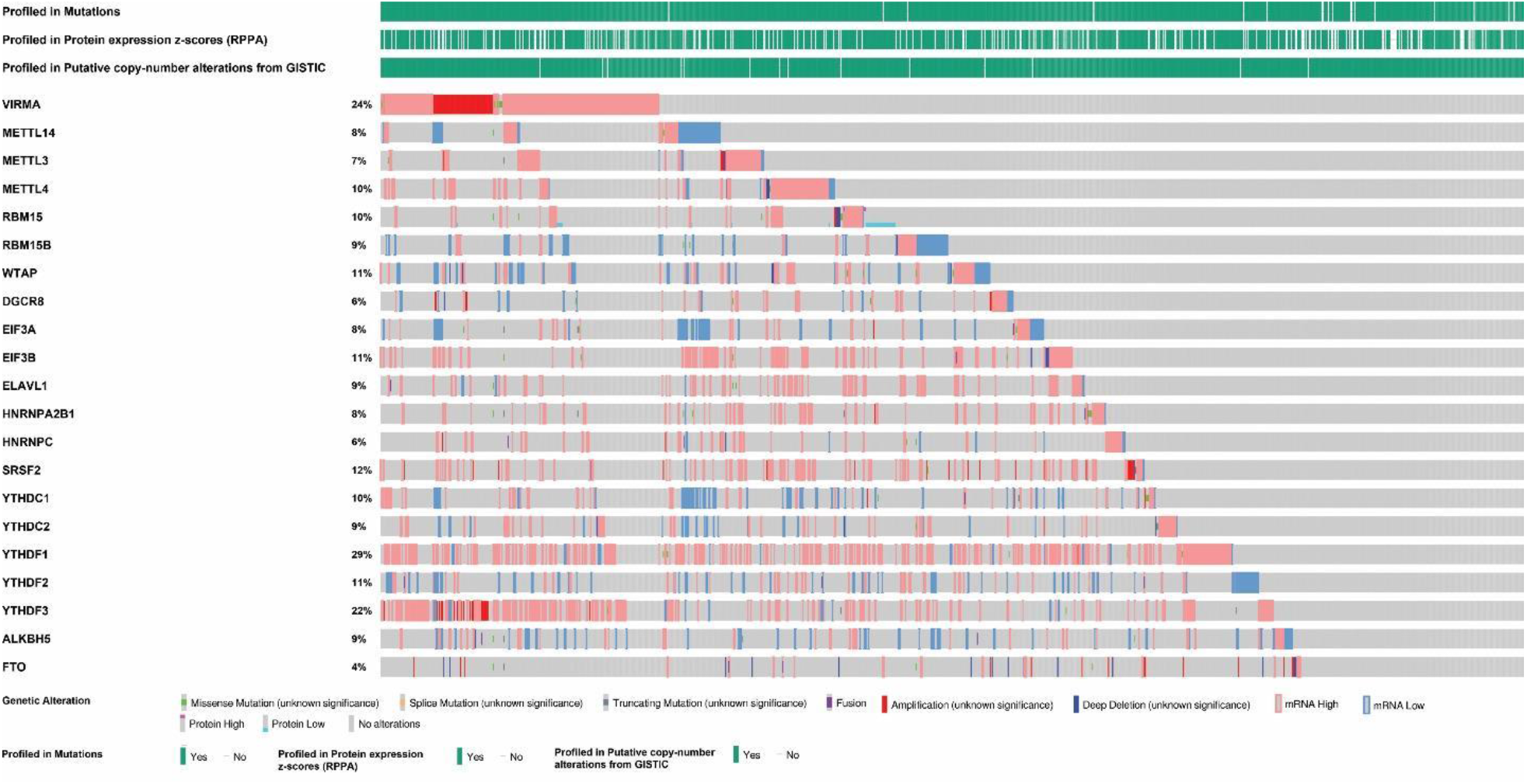
Mutation analysis of m^6^A RNA modification regulatory proteins in breast invasive carcinoma. Oncoprint visualization of TCGA datasets for the m^6^A RNA modifying genes using the cBioportal database. The gene alteration frequencies of m^6^A regulators in BIC were 29% in YTHDF1, 24% in VIRMA, and 22% in YTHDF3, etc.

The frequency of genetic alteration of m^6^A pathway genes (classified as writers: VIRMA, METTL14, METTL3, METTL4, RBM15, RBM15B, WTAP, readers: DGCR8, EIF3A, EIF3B, ELAVL1, HNRNPA2B1, HNRNPC, SFRS2, YTHDC1, YTHDC2, YTHDF1, YTHDF2, YTHDF3, and erasers: ALKBH5, FTO) varies from 4% to 29% (Figure 1). Many types of mutations in m^6^A RNA methylation regulators were observed and particularly amplification, high mRNA levels, and deep deletion were the most abundant genetic alterations for the analyzed genes (Figure 1). Overall, about 80.4% of the samples (820/1084) had genetic changes in at least one of the m^6^A regulators (Figure 2). Among the others, YTHDF1 (29%), VIRMA (24%), and YTHDF3 (22%) genes were found as frequently altered, and FTO (4%), HNRNPC (6%), and DGCR8 (6%) were identified as least altered genes. Remarkably, for frequently altered genes, high mRNA expression (22.8%, 13.6% and 15.2%, respectively), amplification (0.2%, 5.3% and 2.4%, respectively) and multiple alterations (4.0%, 4.8% and 3.4%, respectively) were the commonly detected genetic alterations. These results suggest that m^6^A pathway regulators are frequently mutated in breast cancer and may play an important role in disease pathogenesis.

**Figure 2.**
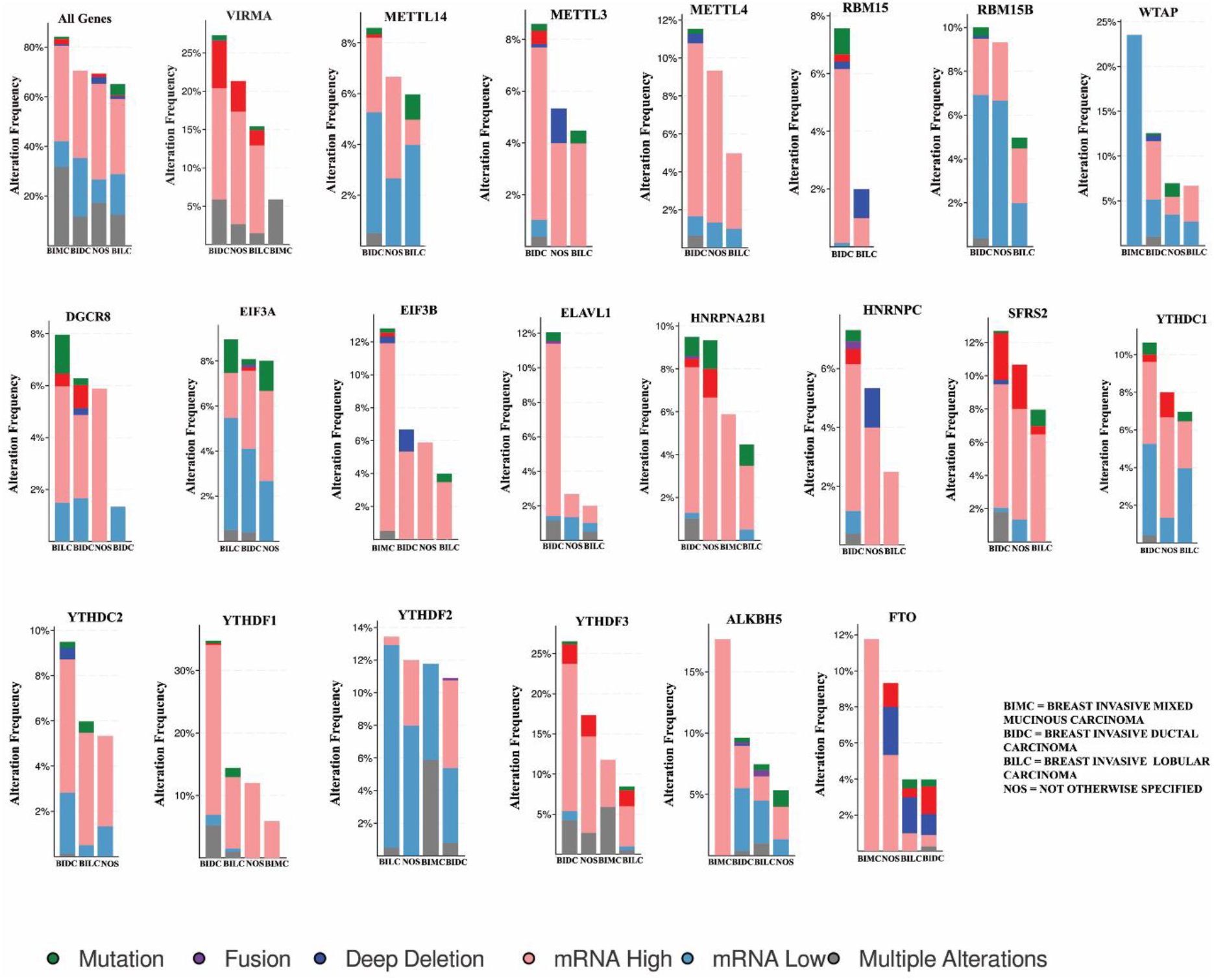
Mutations of m^6^A pathway components in subtypes of breast invasive carcinoma. Mutation, fusion, deep deletion, mRNA high, mRNA low, and multiple alterations were the main genetic alterations and alteration frequency for each gene in subtypes of BIC (Breast Invasive Ductal Carcinoma, Breast Invasive Lobular Carcinoma, Breast Invasive Mixed Mucinous Carcinoma and Not Otherwise Specified) was demonstrated. About 85% of BIMC samples, 70% of BIDC samples, 68% of NOS samples and 64% of BILC samples have at least one of the genetic alterations for m^6^A pathway genes.

To evaluate the distribution of mutation frequency for subtypes of breast invasive cancer, genetic alteration for each gene in Breast Invasive Ductal Carcinoma (BIDC), Breast Invasive Lobular Carcinoma (BILC), Breast Invasive Carcinoma (NOS), and Breast Invasive Mixed Mucinous Carcinoma (BIMC) was investigated (Figure 2). Although for most of the genes (VIRMA, METTL14, METTL3, METTL4, RBM15, RBM15B, ELAVL1, HNRNPA2B1, HNRNPC1, SFRS2, YTHDC1, YTHDC2, YTHDF1, and YTHDF3) genetic alterations were observable in BIDC, BILC turned out to be the main subtype of breast cancer for DGCR8, EIF3A, and YTHDF2 genetic alterations, and the rest of the genes (WTAP, EIF3B, ALKBH5, and FTO) were found as mainly altered in BIMC (Figure 2). The type and frequency of all detected mutations for each gene in all subtypes were shown in detail in Figure 2.

### 3.2 Expresion levels of m^6^A pathway genes revealed significant differences between cancer and control samples

As a complementary analysis to the genetic alteration profile, we next compared the expression levels of m^6^A regulatory genes for primary tumor and control (normal) groups using the TCGA BIC dataset on UALCAN database (http://ualcan.path.uab.edu). Differential expression profile (tumor vs normal) for each gene was illustrated in Figure 3. Expression levels of the 8 genes of the pathway (VIRMA, RBM15, EIF3B, ELAVL1, HNRNPA2B1, HNRNPC, SFRS2, YTHDF1) were significantly upregulated in primary tumor samples, whereas 5 genes (METTL14, WTAP, EIF3A, YTHDC1, FTO) were found as significantly downregulated in primary tumor samples compared to normal ones (Figure 3). No significant difference in the expression level of the other 8 genes (METTL3, METTL4, RBM15B, DGCR8, YTHDC2, YTHDF2, YTHDF3, ALKBH5) was observed between primary tumor and control groups (Figure 3).

**Figure 3:**
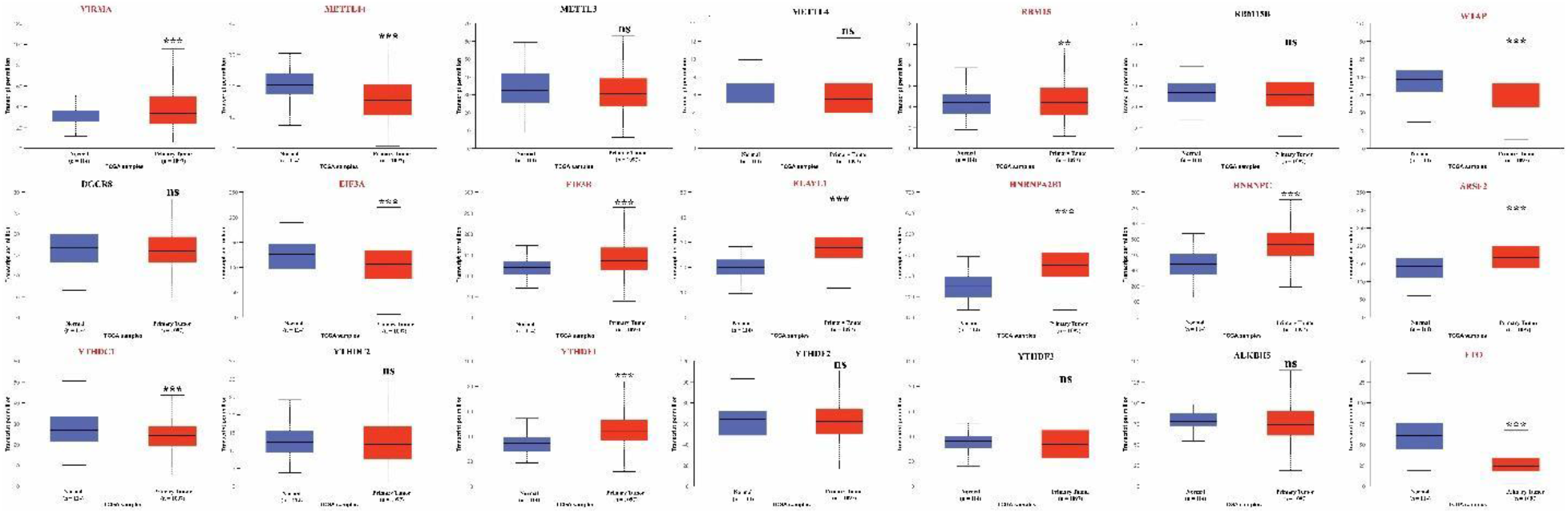
Analysis of expression of m^6^A RNA modification regulatory genes in Breast Invasive Carcinoma. UALCAN dataset was analyzed for the expression of m^6^A genes in 1097 primary tumor samples and 114 corresponding normal tissue samples. Expression of m^6^A genes in patient-derived tumor samples was compared with patient-derived healthy tissue samples. Statistically significant expression levels are shown in red color. ∗*P* < 0.05, ∗∗*P* < 0.01, and ∗∗∗*P* < 0.001 between the primary tumor and normal groups.

To gain more insight into the expression level of m^6^A RNA modifying genes, we then focused on the comparison between pathological stages of cancer (stage I, stage II, stage III, and stage IV) and normal samples (Figure S1 and Table S1). As can be seen from the figure, the trend of the gene expression level in different stages of breast cancer and control samples is highly similar with the comparison based on primary tissue and control samples for most of the investigated genes such as EIF3B, ELVAL1, METTL14, WTAP, METTL3 and METTL4 (Figure S1). Upregulation of EIF3B and ELAVL1 gene expression in primary tissue is also followed in all stages, and for the downregulated genes in primary tissue such as METTL14 and WTAP, under-expression at all stages compared to normal samples was obtained (Figure S1). For the majority of the genes such as METTL3 and METTL4 with no significant differences in expression level between primary tissue and control samples, comparison expression levels between the pathological stages and normal samples was not significant as well (Figure S1). On the other hand, for some genes including VIRMA, EIF3A, and SFRS2, differential expression level between stage IV and control samples was detected as nonsignificant (Figure S1) highlighting the importance of investigation at substages along with the primary tissue level analysis.

### 3.3 Promoter methylation levels poorly overlapped with expression levels of the m^6^A pathway genes

We reasoned that transcriptional alterations of the m^6^A pathway genes could be due to changes at the promoter methylation level. To evaluate this, the significance of methylation level differences between cancer and normal tissue samples for the same gene set was investigated. Previous studies [56] defined the beta value cut-off for hyper-methylation as beta value > 0.7 and for hypo-methylation as beta value <0.3. Remarkably, by employing this cut-off, promoters of all m^6^A pathway genes can be considered as hypo-methylated in both tumor and non-tumor samples except the promoter of DGCR whose beta value is over 0.7 in cancer and normal samples (Figure S2). Differential methylation analyses revealed that promoter of the 7 genes (METTL14, RBM15B, DGCR8, YTHDC1, YTHDF1, YTHDF2, and HNRNPA2B1) were significantly more methylated in tumor samples than the normal ones (Figure S2). Compared with the non-tumor samples, 6 of m^6^A pathway genes (METTL3, METTL4, EIF3B, YTHDC2, SRSF2, and HNRNPC) in tumor tissues were found as significantly less methylated, and there was no significant difference at methylation level of the promoter for the rest of the genes (Figure S2). Loss of the correlation between promoter methylation and expression levels was notable.

Next, we checked the methylation status of the promoters for four pathological stages of breast invasive cancer (Figure S3). The result demonstrated that the methylation level of promoters for pathological stages was generally in accordance with the trend of the methylation profile shown in Figure S2. However, as in the expression level, the significance of methylation alteration between stage IV and control samples was identified contradictory to the general trend (Figure S3), which emphasized the necessity of analysis at stage level complementary to primary tissue level investigation. Significant alterations were marked in the figures and associated p-values are listed in Table S2.

### 3.4 Association of expression level and mutation frequency of m^6^A pathway genes and the prognosis of cancer patients

Clinical information in the TCGA database was retrieved and the overall survival metric was considered to explore the influence of the pathway genes expression on survival rate. The patient’s survival rate for high and low expression of the m^6^A genes was evaluated through the use of the Kaplan-Meier plotter (https://kmplot.com/) [51] (Figure 4). Of these genes, 6 of them have a significant effect on the survival rate. High expression of VIRMA, METTL14, and RBM15B genes was significantly associated with longer overall survival in patients (p-value <0.05, Figure 4). Noticeably, low expression of EIF3B, YTHDF1, and YTHDF3 was associated with significantly high overall survival compared to the patients with high gene expression levels (Figure 4). Overall survival of the patients was similar for the rest of the genes regardless of the expression levels of the genes. This analysis extrapolated the association of the prolonged survival and expression levels of several m^6^A pathway components in breast invasive carcinoma.

**Figure 4.**
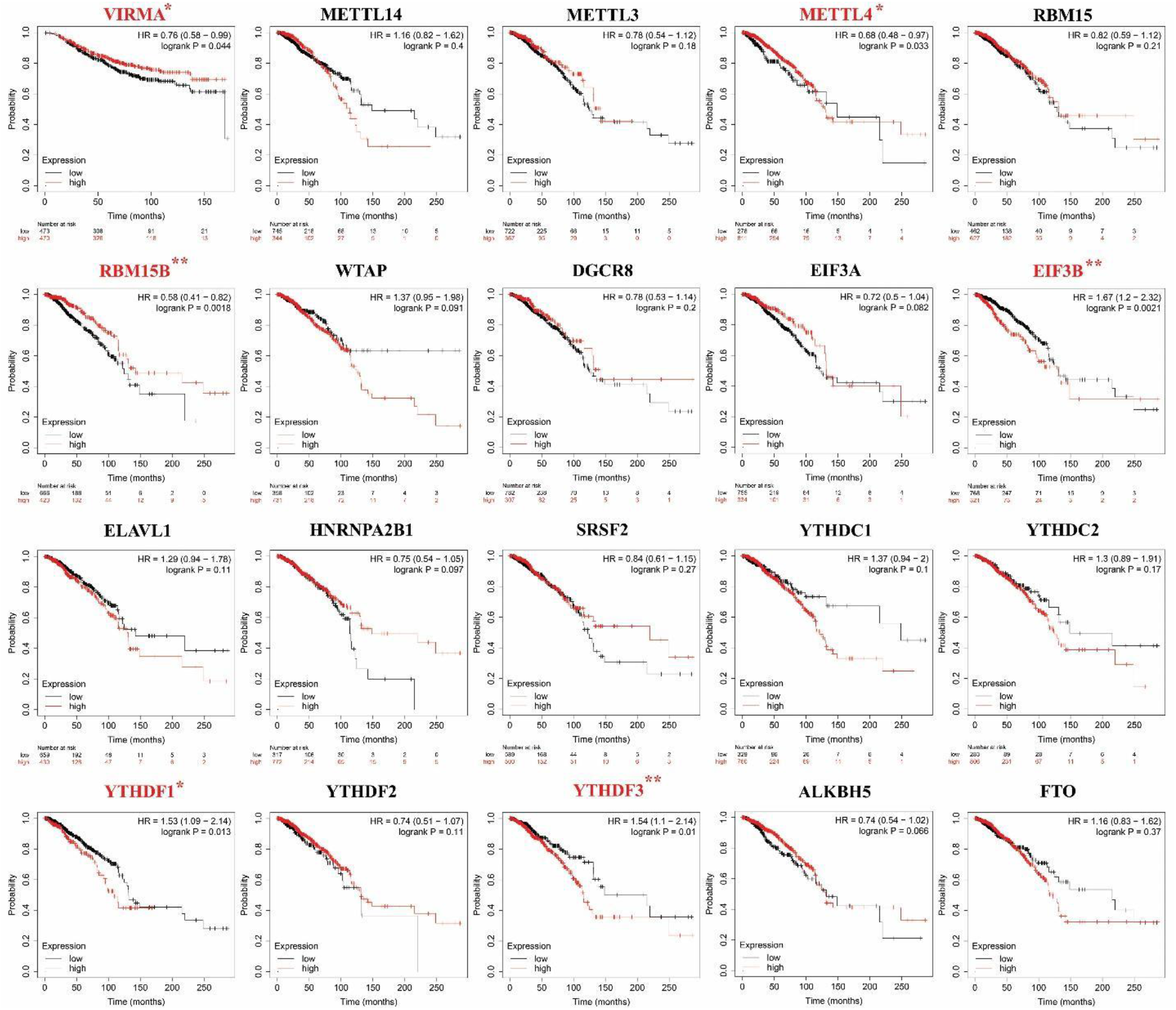
Kaplan–Meier survival curves of m^6^A regulatory genes in breast invasive carcinoma regarding the expression level. Overall survival curves were calculated and drawn based on the differential expression levels of m^6^A regulatory genes in BIC. The black and red curves represented the survival curves of the wild type and all mutation groups, respectively. Statistically significant expression levels are shown in red. ∗*P* < 0.05 and ∗∗*P* < 0.01 between the two groups.

Furthermore, the prognostic value of mutation frequency of examined genes in breast cancer was assessed. A significant association between breast cancer patients’ survival and mutation frequency of the m^6^A pathway genes was sought and specifically, patients with tumors displaying the mutations in one of the nine genes (METTL14, RBM15, WTAP, HNRNPA2B1, SRSF2, YTHDF1, YTHDF2, YTHDF3, and ALKBH5) had significantly lower overall survival in comparison to mutation free patients (Figure S4).

### 3.5 Analysis of m^6^A pathway proteins’ expression in cancer and normal tissue

To further dissect the potential link between the expression level of m^6^A genes and breast invasive cancer, protein expression profile in cancer and normal tissues was inspected using The Human Protein Atlas data [52]. Human Protein Atlas is a valuable online database with the spatial expression data of proteins, and its broad antibody collection allows us to map and compare the proteins in tumors and corresponding normal tissues. Among our list, 3 of the investigated genes (METTL3, YTHDF1, and YTHDF3) have not been profiled in the atlas yet. Based on the scoring system of the database, 1 of the genes were identified as lowly expressed (<25%), 2 of them were considered as moderately expressed (25% to 75%) and 15 of them were found as highly expressed (>75%) (Figure 5). IHC staining of m^6^A genes coding proteins for the breast invasive cancer and corresponding normal counterpart tissues confirmed that these proteins were overexpressed in BIC samples compared to normal tissue (Figure 5). According to the IHC data, there is a poor correlation of upregulated mRNA levels and increased protein levels for m^6^A pathway genes.

**Figure 5.**
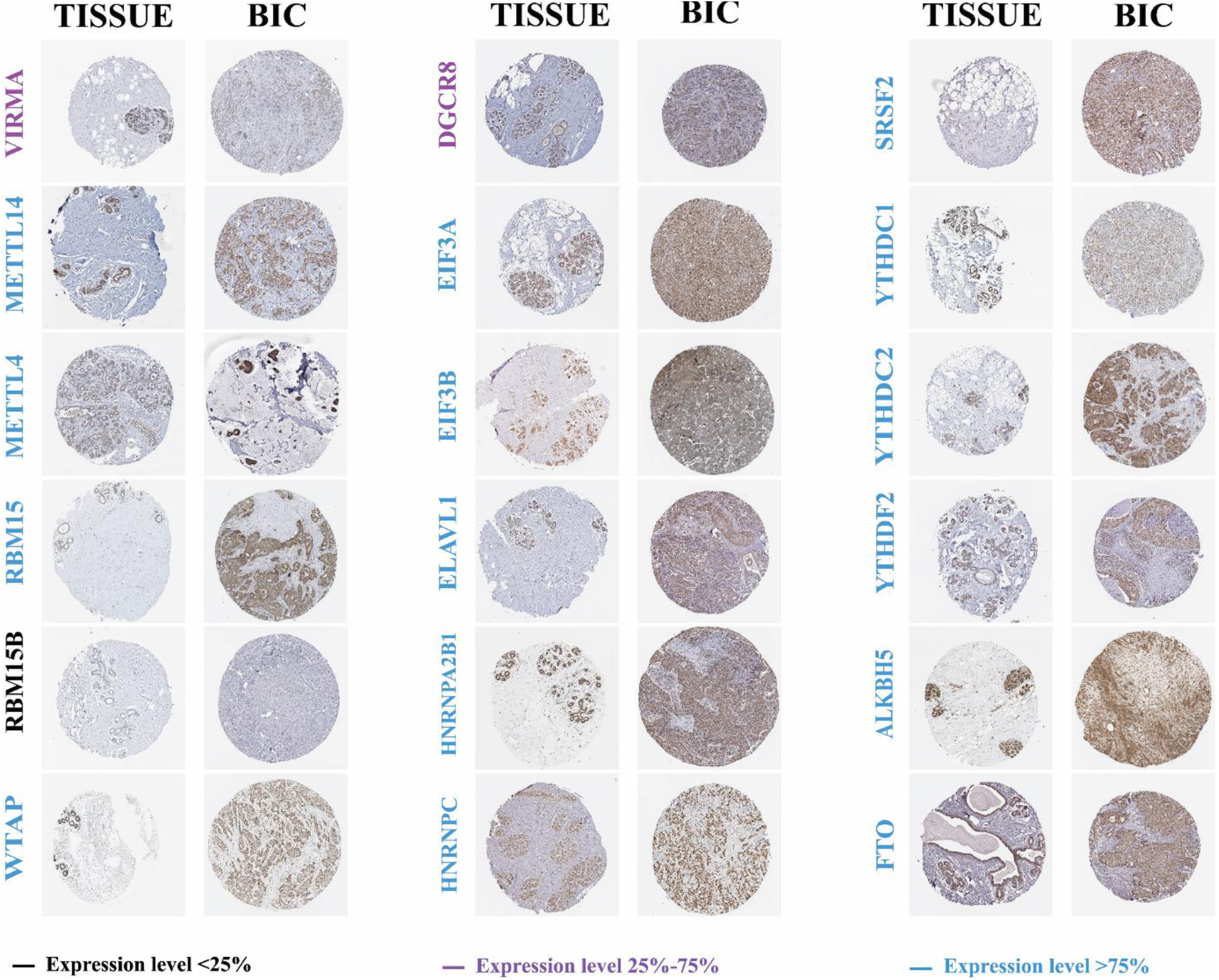
Analysis of expression of m^6^A regulatory proteins in Breast Invasive Carcinoma through the use of The Human Protein Atlas. Representative images of immunohistochemical staining of the m^6^A regulatory genes in patient-derived breast cancer samples as well as healthy breast tissue samples from The Human Protein Atlas data.

### 3.6 *Enriched* pathways associated with genetic alterations in the m^*6*^A *p*athway

We then analyzed the transcripts whose levels were modulated upon genetic alteration (mutation, amplification, deep deletion, high or low mRNA levels) of m^6^A regulatory genes. The altered group consists of the patients’ samples with genetic alteration in the m^6^A pathway and the unaltered group is formed by the patients’ samples without genetic alteration. DEGs (p<0.01 and |FC|>1.5) between altered (310 genes) and unaltered (1186 genes) groups were displayed on a volcano plot (Figure 6A) and listed in Table S3. The selected top 10 DEGs for altered (PRAME, A2ML1, MYBL2, INAVA, BIRC5, MCM10, UBE2C, CBX2, PLCH1, and ONECUT2) and unaltered (ANKRD30A, PIP, TFF1, SCGB2A2, ADH1B, PTPRT, CAPN8, HMGCS2, NEK10, and LRRC31) were highlighted on the volcano map (Figure 6A). In order to explore the GO terms enriched by the DEGs, firstly, a merged list of altered and unaltered DEGs was enriched to depict the terms at BP, MF, and CC levels. All the identified terms were summarized in Table S4. Top 5 BPs (‘organelle fission’, ‘nuclear division’, ‘mitotic nuclear division’, ‘mitotic sister chromatid segregation’ and ‘sister chromatid segregation’) formed two main clusters with enriched altered and unaltered genes were displayed in a cnet figure (Figure 6B). The presence of shared DEGs in different pathways within the clusters is notable and these DEGs might take a role in cross-talking of the pathways.

**Figure 6.**
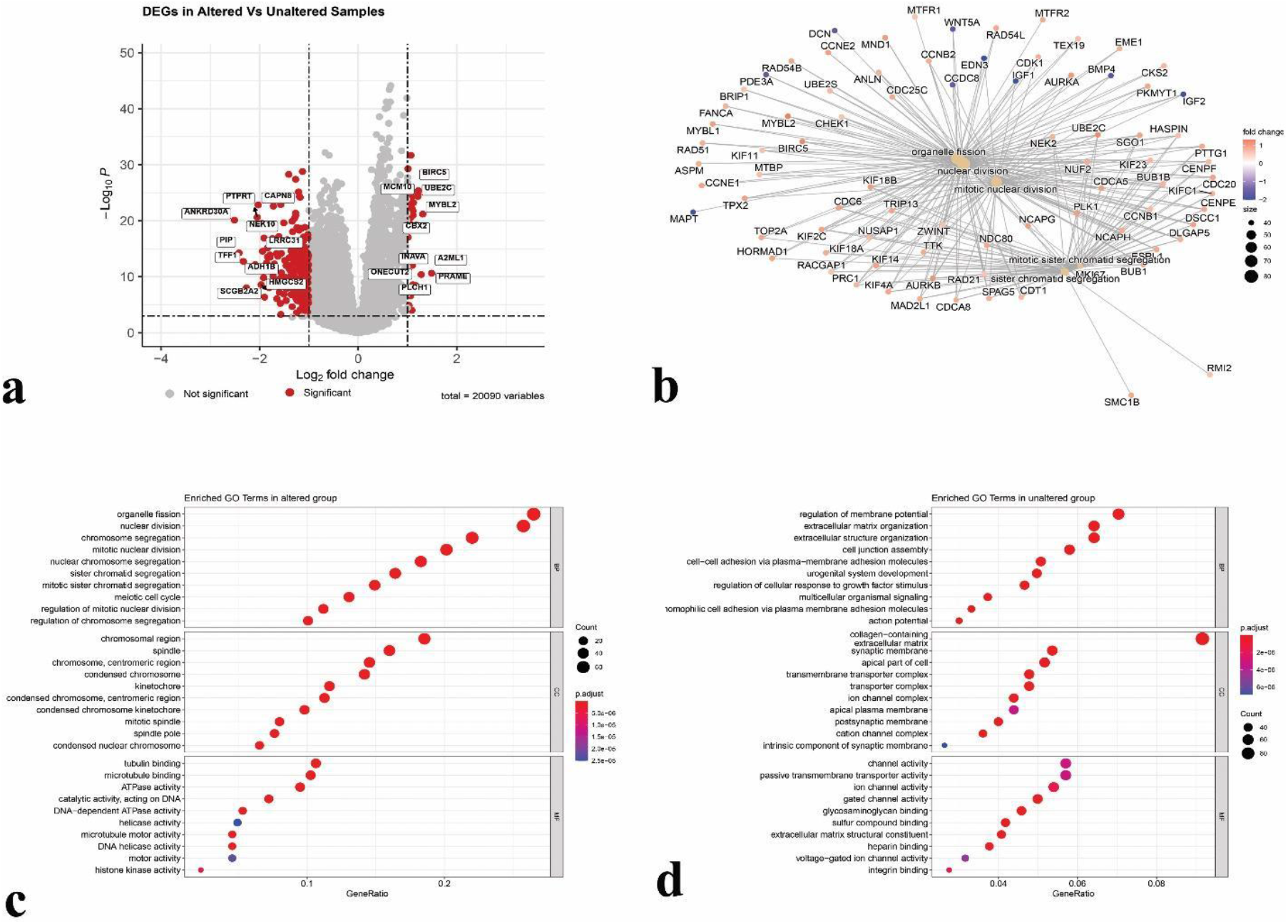
Differential expression of genes and overrepresented pathways. a) Volcano plot depicting the significance of the genes (p < 0.05) in altered and unaltered groups. b) Gene-concept network of the top 5 overrepresented pathways enriched by the DEGs in altered and unaltered lists. Dotplot of the top 10 enriched terms of each GO category (BP, MF, and CC) by the DEGs in C) m^6^A altered group (upregulated genes as a result of genetic alteration in m^6^A pathway) D) m^6^A unaltered group (downregulated genes as a result of genetic alteration in m^6^A pathway)

As the following analysis, DEGs in altered and unaltered lists were enriched separately to figure out the pathways associated with altered and unaltered samples resulting in significant (p<0.05) 243 BPs, 17 MFs and 39 CCs for altered, and 373 BPs, 51 MFs and 49 CCs for the unaltered group which was summarized in Table S5. Dot plots were constructed using the enriched GO terms for both groups (Figure 6C-D). Surprisingly, almost all of the top 10 detected BPs (‘nuclear division’, ‘chromosome segregation’, ‘mitotic nuclear division’, ‘sister chromatid segregation’, ‘nuclear chromosome segregation’, ‘mitotic sister chromatid segregation’, ‘regulation of chromosome segregation’, ‘regulation of mitotic nuclear division’ and ‘meiotic cell cycle’) in altered groups (Figure 6C) were strongly associated with proliferation which may indicate that genetic alteration in m^6^A pathway components had a remarkable impact on cell division. Those DEGs in the altered list enriched several cellular components, such as ‘chromosomal region’, ‘spindle’, ‘centromeric region’, ‘condensed chromosome’, and ‘kinetochore’ (Figure 6C). The genes are also enriched in various molecular functions such as ‘tubulin binding’, ‘microtubule binding’, ‘ATPase activity’, ‘catalytic activity acting on DNA’, and ‘DNA-dependent ATPase activity’ (Fig 6C). Enriched terms in MF and CC categories provided support to the potential effect of genetic alteration in m^6^A genes on cell cycle-related biological processes.

Same enrichment was conducted using the DEGs list for unaltered samples and the top terms in all categories were shown in figure 6D. BPs such as ‘regulation of membrane potential’, ‘extracellular matrix organization’, ‘extracellular structure organization’, ‘cell junction assembly’, and ‘cell-cell adhesion via plasma-membrane adhesion molecules’ were found among the top 10 BPs enriched by the genes in unaltered samples (Figure 6D). ‘Collagen containing extracellular matrix’, ‘synaptic membrane’ and ‘apical part of the cell’ were identified as top CC terms, and ‘channel activity’, ‘passive transmembrane transporter activity’ and ‘ion channel activity’ were the top MF terms (Figure 6D).

An alternative method to the over-representation test is an application of one of the functional class scoring approaches such as gene set enrichment analysis (GSEA). To further extend the information on biological processes we employed GSEA using the same set of significant mRNAs without a fold change cut-off. The appearing list of genes was enriched in all ontologies (BP, MF, and CC) and KEGG pathways (Figure S5 and Table S6). Aside from cell cycle-related biological processes (i.e. ‘cell cycle’, ‘mitotic cell cycle’, ‘’mitotic cell cycle process’, ‘nuclear division’), RNA binding and processing related processes such as ‘RNA binding’, ‘ncRNA metabolic process’, ‘ncRNA processing’, ‘mRNA processing’ and organelle organization related process such as ‘ribosome biogenesis’ were enriched for the over-expressed genes in altered samples (Figure S5A). Whereas the top BPs enriched by over-expressed genes in unaltered samples were found as ‘extracellular matrix’, ‘collagen-containing extracellular matrix’, ‘biological adhesion’, and ‘cell adhesion’ (Figure S5A). Moreover, enriched KEGG pathways by over-expressed mRNAs in altered and unaltered samples were portrayed on a ridge plot (Figure S5B) and listed in Table S6. Several pathways, as ‘spliceosome’, ‘RNA transport’ and ‘cell cycle’, were enriched by the altered samples and over-expressed genes in unaltered samples enriched the pathways such as ‘cell adhesion molecules’ and ‘ECM-receptor interaction’ (Figure S5B).

### 3.7 Interaction of downstream pathway proteins

Next, we interrogated the interaction of differential proteins in top BP terms with m^6^A pathway regulatory proteins via the implementation of the STRING database [53]. By being the most significant terms, ‘nuclear division’ for the altered group with 72 differential proteins and ‘regulation of membrane potential’ for the unaltered group with 70 differential proteins were prioritized to carry out this analysis. The protein-protein interaction network was mapped (Figure 7) and UBE2C, CCNE1, CCNE2, and PKMYT1 were identified as important connections between nuclear division network and m^6^A pathway proteins (Fig 7A). For the interaction of regulation of membrane potential and m^6^A pathway proteins, GLRB and MAPT were found as chief connections (Figure 7B).

**Figure 7.**
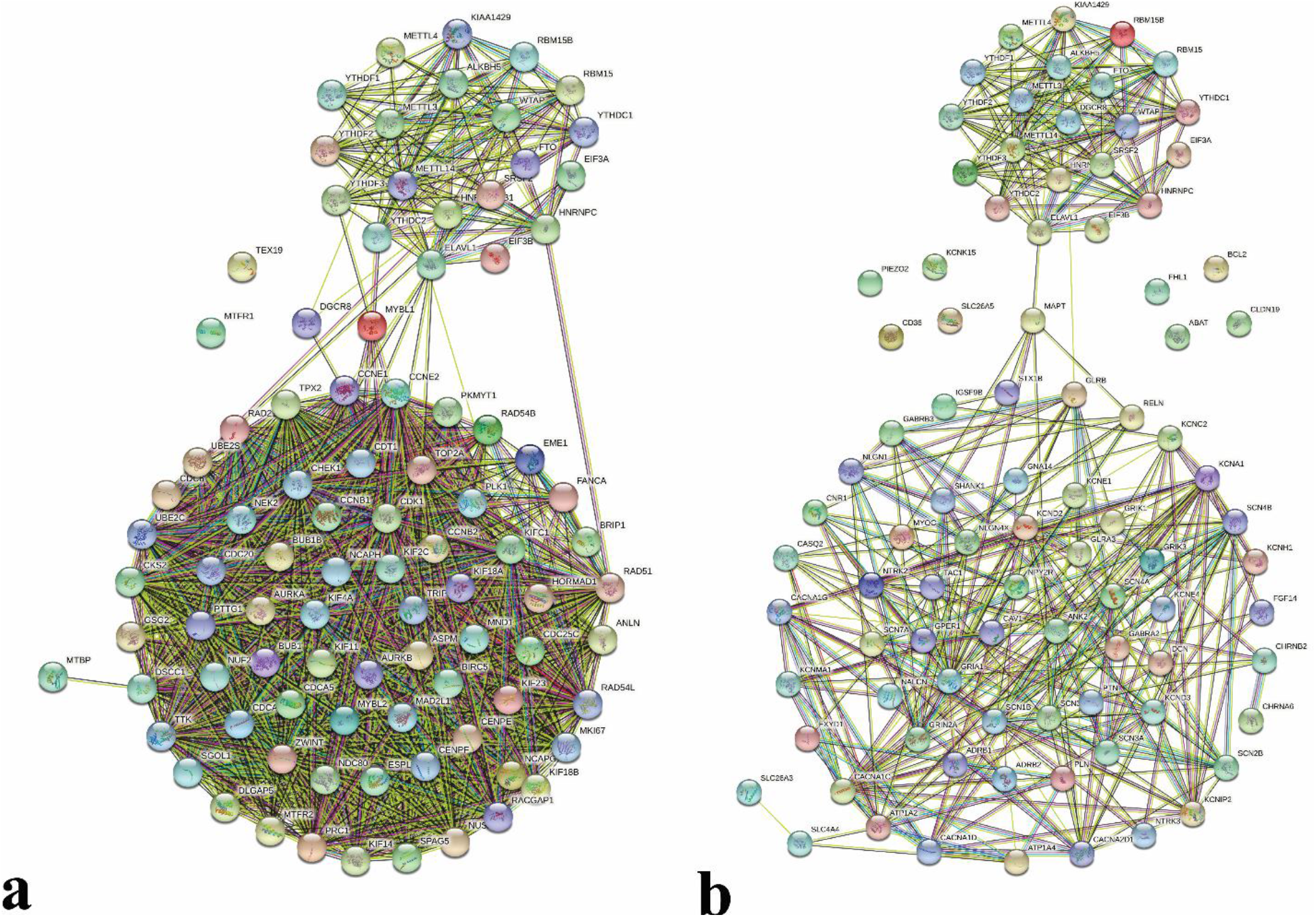
Multicenter protein-protein interaction (PPI) network of top significant BPs and m^6^A pathway. PPI network was constructed between m^6^A regulatory proteins and differential proteins in the (A) Nuclear division and (B) regulation of membrane potential pathways using the STRING database.

### 3.8 Enriched pathways associated with genetic alterations in m^5^C and m^1^A pathways

The compelling link established by our analyses between genetic alterations in m^6^A genes and cancer-associated pathways such as cell cycle, DNA repair, and RNA biogenesis processes promoted us to inspect the m^5^C and m^1^A pathways whose components take roles in methylation of cytosine and adenosine of RNA, respectively. The altered gene expression profile upon genetic alteration in m^5^C pathway genes was retrieved and analyzed as did for m^6^A pathway components, yielded 646 DEGs in altered and 796 DEGs in unaltered samples (p<0.01 and |FC|>1.5). The same examination was conducted for m^1^A regulator mutation as well and 252 and 345 DEGs were identified in altered and unaltered samples, respectively. GO terms over-representation analyses of the DEGs in altered and unaltered groups for m^5^C and m^1^A were performed, presented in Table S7, and displayed on Figure 8. Very remarkably, top enriched BPs by upregulated genes such as ‘chromosome segregation’, ‘nuclear division’, ‘mitotic nuclear division’, ‘sister chromatid segregation’ and ‘mitotic sister chromatid segregation’ in both genetically altered samples with genetic alteration in m^5^C and m^1^A pathway genes were related to cell cycle and proliferation as observed in altered list of m^6^A regulatory components (Fig 8A, C). DEGs in altered samples in both m^5^C and m^1^A lists exhibited similar CC terms such as ‘chromosome, centromeric region’, ‘condensed chromosome, centromeric region’ and ‘spindle’ (Fig 8A, C). MF terms such as ‘microtubule binding’, ‘chemokine activity’, and ‘tubulin binding’ were common for both lists. These results highlighted the strong correlation of the enriched terms for the upregulated DEGs in the patient samples with genetic alterations in m^5^C and m^1^A pathway genes.

**Figure 8.**
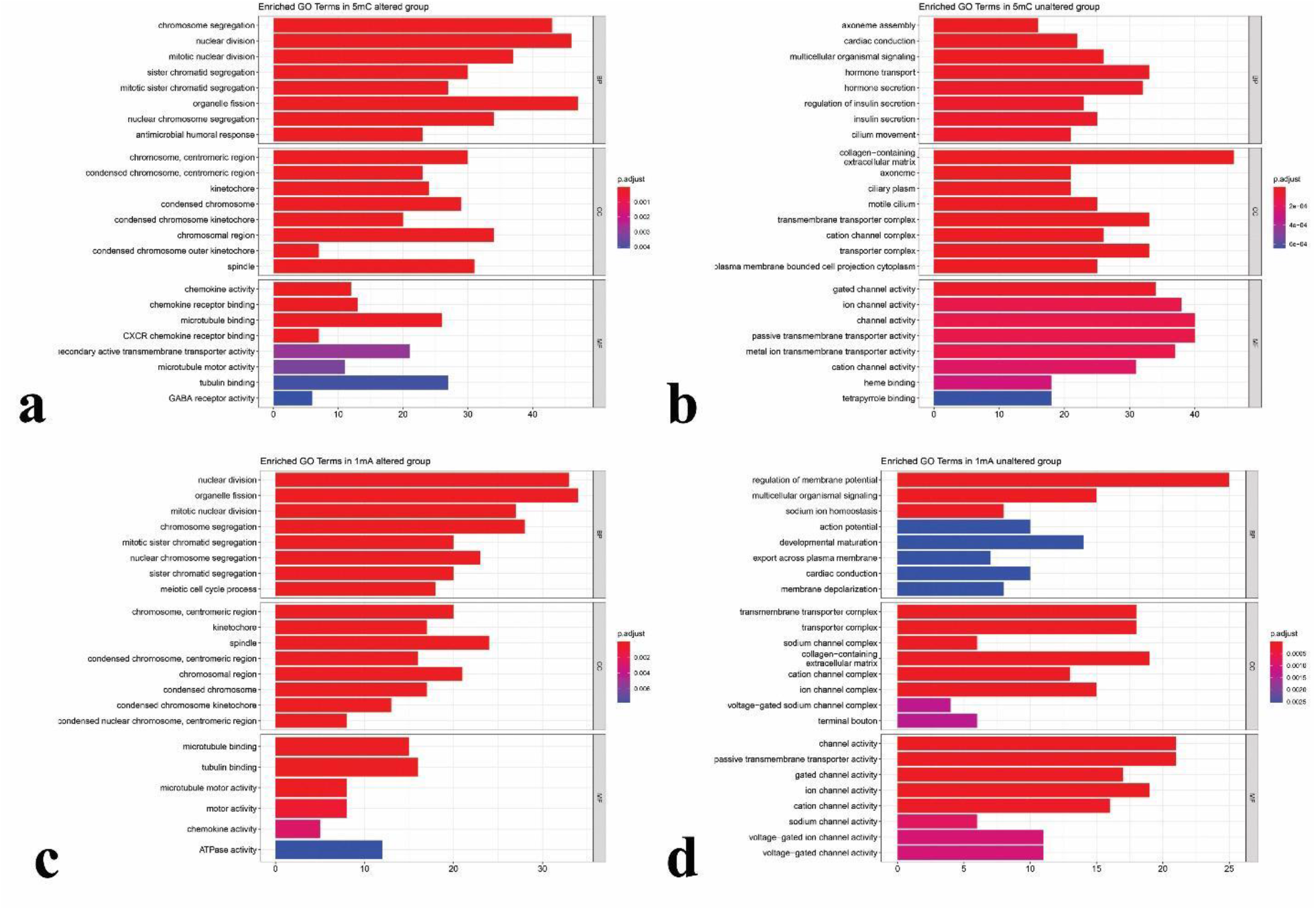
GO of the differentially expressed genes in altered and unaltered groups. Barplots illustrating the top enriched terms of each GO category (BP, MF, and CC) by the DEGs in: A) m^5^C altered group (upregulated genes as a result of genetic alteration in m^5^C pathway) B) m^5^C unaltered group (downregulated genes as a result of genetic alteration in m^5^C pathway) C) m^1^A altered group (upregulated genes as a result of genetic alteration in m^1^A pathway) D) m^5^C unaltered group (downregulated genes as a result of genetic alteration in m^1^A pathway)

Top 5 enriched terms by the upregulated DEGs in the patient samples without genetic alterations in m^5^C pathway genes were ‘axoneme assembly’, ‘cardiac conduction’, ‘multicellular organismal signaling’, ‘hormone transport’ and ‘hormone secretion’ at BP level, ‘collagen-containing extracellular matrix’, ‘axoneme’, ‘ciliary plasm’, ‘motile cilium’ and ‘transmembrane transporter complex’ at CC level, and ‘gated channel activity’, ‘ion channel activity’, ‘channel activity’, ‘passive membrane transport activity’ and ‘metal ion transmembrane transport activity’ at MF level (Figure 8B). Several terms were enriched by the upregulated DEGs in the patient samples without genetic alterations in m^1^A pathway genes, and the top 5 terms were as following: ‘regulation of membrane potential’, ‘multicellular organismal signaling’, ‘sodium ion homeostasis’, ‘action potential’ and ‘developmental maturation’ for BPs, ‘transmembrane transporter complex’, ‘transporter complex’, ‘sodium channel complex’, ‘collagen-containing extracellular complex’ and ‘cation channel complex’ for CCs, ‘channel activity’, ‘passive membrane transport activity’, ‘gated channel activity’, ‘ion channel activity’ and ‘cation channel activity’ for MFs (Figure 8D).

### 3.9 Common downstream pathways for m^*6*^A, m^*5*^C and m^1^A genetic alterations

Overlapping top BPs enriched by upregulated DEGs in the patient samples with genetic alterations in any of the m^6^A, m^5^C or m^1^A pathways was noteworthy, and this observation stimulated us to monitor the common and unique BPs among these pathways. The distribution of the enriched BPs among the three pathways was depicted on a Venn diagram, and 42 common BPs were identified (Figure 9A). The number of unique BPs (175 in total, in m^6^A 55 BPs and m^5^C 120 BPs) was not far greater than shared BPs, indicating the notion of common BPs as evidence of a similar impact of these three pathways in breast invasive carcinoma. 42 common BPs were displayed on a heatmap (Figure 9B) and proliferation-related BPs such as ‘nuclear division’, ‘chromosome segregation’, and ‘mitotic nuclear division’ were evident for the identified common BPs.

**Figure 9.**
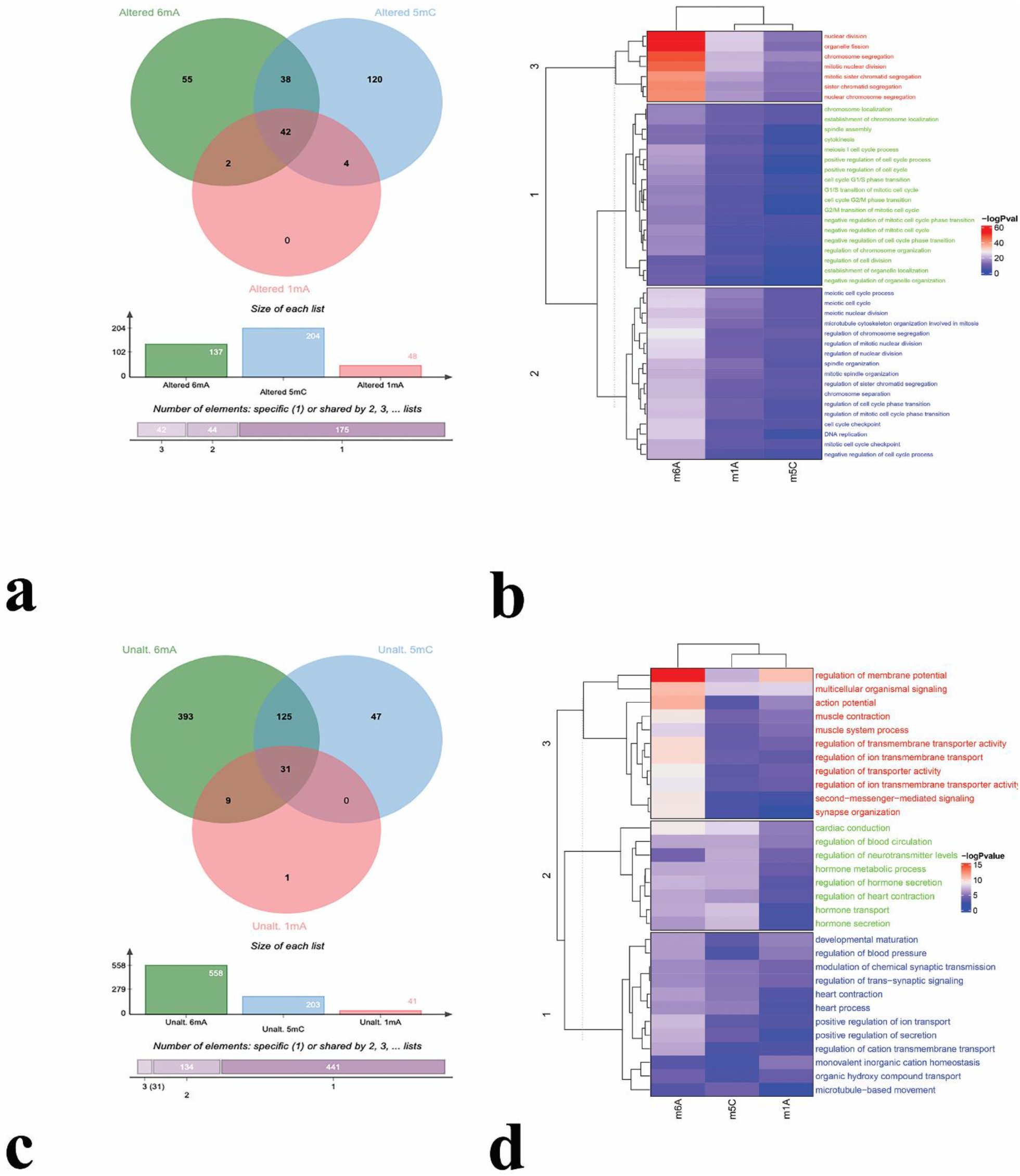
Comparison of differentially expressed genes in altered and unaltered groups of epitranscriptomic pathways. Venn diagrams portraying the commonality between or uniqueness of DEGs a) in altered and b) in unaltered groups of m^6^A, m^5^C, and m^1^A pathways. Clustered heatmap of common BPs enriched by the DEGs of C) altered and D) unaltered groups. −logp value is visualized in color scale from purple to red as an indicator of the significance level.

For the upregulated DEGs in the patient samples without genetic alterations in any of the m^6^A, m^5^C, or m^1^A pathway genes, the same analysis resulted in 31 common and 441 unique (393 for m^6^A, 47 for m^5^C, and 1 for m^1^A) BPs (Figure 9C). As shown in Figure 9D, common BPs were not closely related. Diverse BPs such as ‘regulation of membrane potential’, ‘multicellular organismal signaling’, and ‘muscle contraction’ specified the top BPs that are commonly observed among three comparisons. Common and unique DEGs among three comparisons were also identified and the obtained results were demonstrated in Figure S6.

## 4. Discussion

More than 170 different types of RNA modifications have been characterized so far, and among these modifications, several of them affect numerous biological processes by acting on mRNA [13]. Chemical modification of mRNA is highly diverse and its effect is considered to be highly specific to regulate the stability, translation efficiency, translocation, and splicing of the mRNAs [57,58]. Particularly, *N*6-methyladenosine (m^6^A) is the most abundant mRNA modification, and m^6^A is co-transcriptionally generated by a methyltransferase complex [58,59]. This study presents a comprehensive analysis of the impact of genetic alteration in m^6^A pathway genes on breast invasive carcinoma.

In this study, we first explored the genetic alteration frequency of m^6^A pathway genes in BIC. YTHDF1 (29%), VIRMA (24%), and YTHDF3 (22%) were found as the highly altered genes in BIC, and mRNA level differences were the dominant type of genetic alterations (Figure 1). YTH N6-methyladenosine RNA binding proteins 1, 2, and 3 (YTHDF1, YTHDF2, and YTHDF3; respectively) are members of the YTH domain family and act as reader proteins of m^6^A modification. YTHDF1 promotes and facilitates translation initiation and increases the translation efficiency [26], whereas YTHDF2 negatively regulates the translation of m^6^A-modified mRNAs by functioning in the decay of methylated mRNAs [27]. Dual roles of YTHDF3 in mRNA stability and translation are manifested in elevated protein synthesis via interaction with YTHDF1 and degradation of methylated mRNAs through YTHDF2 [60]. Over-expression and activity of YTHDF1 are associated with hepatocellular carcinoma [61], melanoma and colon cancer [62], and colorectal cancer [63,64] by modulating the cell proliferation, stemness, and anti-tumor immunity processes. A recent study demonstrated the over-expression of YTHDF1 and YTHDF3 is due to frequent amplification of YTHDF1 and YTHDF3 in breast cancer and is linked to metastasis and poor prognosis in breast invasive cancer [65]. According to this study, the synergetic interaction between YTHDF3 and YTHDF1 proteins enhances the translation efficiency of methylated mRNAs and this might be the key step to regulate the tumorigenicity [65]. VIRMA (Vir-like m^6^A methyltransferase associated protein) is one of the writer proteins functioning in the methylation of mRNA and non-coding RNAs [66]. Upregulated VIRMA levels in multiple cancers such as lung squamous cell carcinoma (LUSC), lung adenocarcinoma (LUAD), liver hepatocellular carcinoma (LIHC), stomach adenocarcinoma (STAD), and breast invasive carcinoma (BRCA) have been demonstrated in a study covering the pan-cancer datasets [67]. According to current literature, increased levels of m^6^A modification and its impact on diverse biological processes by the activity of reader proteins are essential in carcinogenesis and our data on genetic alterations in m^6^A pathway components confirmed this notion for breast invasive cancer.

Across different subtypes of breast cancer, the most commonly observed one is Breast Invasive Ductal Carcinoma (BIDC) with 70%–75% of the cases, followed by Breast Invasive Lobular Carcinoma (BILC) with 5%–15% of the cases [68,69]. While mapping genetic alterations in breast invasive cancer recognize all samples coming from different subtypes, subsequent analyses highlighted that alterations are concentrated mainly in BIDC followed by BILC (Figure 2). BILC has a better prognosis in comparison to BIDC with a low histological grade and a good response to hormone therapy [70]. Since genetic alterations in m^6^A pathway components are generally associated with the progression and aggressiveness of cancer [32,71,72] our obtained results on the accumulation of genetic alterations of this pathway members in BIDC cases rather than BILC ones are consistent with the previously published studies.

We noticed that the expression level of several m^6^A pathway genes was altered between BIC and normal tissues (Figure 3). This finding agreed well with data from the in-depth gene expression analyses at pathological stages (Figure S1). However, inconsistency in expression level data at stage IV underlined the necessity of investigation at substages along with the primary tissue level for diagnostic or prognostic use of these genes as biomarkers. Significant overexpression of VIRMA, RBM15, EIF3B, ELAVL1, HNRNPA2B1, HNRNPC, SFRS2, and YTHDF1, and downregulation of METTL14, WTAP, EIF3A, YTHDC1, and FTO in BIC samples compared to normal tissues were observed. It was noteworthy to observe that some of the writer and reader proteins were significantly upregulated and a couple of them were significantly downregulated in cancer samples compared to normal tissues. The opposite trend in expression levels for the genes with similar catalytic activity might be due to the specificity of the interactions between m^6^A components and target mRNAs or redundancy of the proteins for functional compensation of the m^6^A pathway. Further studies are needed to unveil the molecular causes of opposite expression directionality. Furthermore, expression level and function variation of m^6^A genes among the cancer types may imply the tumor type-specific roles of this pathway. As an example, METTL14 can act as a tumor-suppressor as in colorectal cancer [73], renal cell carcinoma [74] and breast cancer [75], or oncogene as in pancreatic cancer [76] which might be due to the target RNA composition specific to cancer types. Therefore, as performed in this study, analysis of these genes using the comprehensive datasets would be more informative to approximate the exact function of the genes across the possible multiple roles of m^6^A pathway components in investigated cancers.

Regarding the genomic instability in cancer cells, gene expression level shift in cancer might be related to copy number alterations (CNA) as shown in previous studies [77,78]. Our data revealed that CNA is not the distinguished source of genetic alterations in m^6^A pathway components and not plausible enough to explain the changes at mRNA levels in breast cancer patients. Therefore, promoter methylation levels were examined to elucidate the molecular mechanism of altered gene expression levels with negligible CNA events (Figure S2-3). Hypo-methylation of the m^6^A pathway genes’ promoters in both cancer and normal tissues pinpointed the active transcriptional methylation signature of these genes for both cancer and normal tissues. Limited overlap of the genes with significant differences at promoter methylation status and gene expression level signified that gene expression level is regulated most probably by other mechanisms such as chromatin organization rather than promoter methylation status.

The expression of m^6^A RNA methylation regulators was tested for their prognostic signature, and 6 of them (VIRMA, METTL14, RBM15B, EIF3B, YTHDF1, and YTHDF3) related to overall survival were identified as predictive genes (*p* < 0.05). Our study indicated that high expression of VIRMA, METTL14, and RBM15B, and low expression of EIF3B, YTHDF1, and YTHDF3 is correlated with longer OS, and these genes could serve as markers to predict the prognosis of BIC (Figure 4). There are common and unique genes in the final set of identified genes as prognostic markers in our and the other cancer studies [63,75,79] which highlight the diverse roles of the m^6^A pathway genes in the progression of different human malignancies. To get more insights into the underlying mechanisms, the association of mutation frequency of the genes with prognostic value was examined and nine of the m^6^A pathway genes (METTL14, RBM15, WTAP, HNRNPA2B1, SRSF2, YTHDF1, YTHDF2, YTHDF3, and ALKBH5) were correlated with the prognosis of BIC (Figure S4). Mutations in these genes are related to lower overall survival in BIC. METTL14, YTHDF1, and YTHDF3 were measured as predictive genes in BIC for both expression level and mutation frequency and may be considered as promising targets in novel therapies of BIC. However, the low number of patients with mutations is a limiting factor to elaborate the putative association between overall survival and the mutation status of the studied genes.

Moreover, we performed an IHC analysis to study the protein expression profile of m^6^A pathway components at the protein level based on The Human Protein Atlas database (Figure 5). Protein expression data of normal and cancer tissues were compared and out of 18 proteins, 15 proteins were highly expressed (>75%) in the majority of the cancer specimens indicating the dysregulation of m^6^A regulatory proteins in BIC. Similar results were previously reported in a bunch of cancers such as melanoma [80], liver cancer [36], and colorectal cancer [81].

In order to shed light on downstream processes affected by genetic alterations in the m^6^A pathway, we sought to explore the significant DEGs and significantly enriched pathways between altered and unaltered groups (Figure 6). The top upregulated genes in the altered group conferring critical roles in proliferation were remarkable. Upregulated ‘preferentially expressed antigen in melanoma’ (PRAME) expression is associated with the progression of several cancers including breast cancer [82], hematological malignancies [83], lung cancer [84], ovarian cancer [85] and melanoma [86]. PRAME is a repressor of retinoic acid receptor-mediated differentiation and apoptosis [87] and increased expression levels of PRAME provide an advantage to breast cancer cells by enhancing cell motility and invasion [88]. MYBL2, another significantly upregulated gene in the altered group, is a member of MYB transcription family and a positive regulator of the cell cycle which binds to the promoter of G2/M phase genes, such as CCNB1, CDK1, CCNA2, BCL2, and BIRC5 [89,90]. Over-expression of MYBL2 is defined in several cancers such as breast [91] lung [92], esophagus [93], colorectal [94], bladder [95] and prostate [96] cancers. BIRC5 is a well-characterized anti-apoptotic gene playing a role in the presentation of apoptosis, and our results described it as one of the top upregulated genes in the altered group compared to the unaltered samples. High expression of *BIRC5* during tumorigenesis in various cancer types including breast [97,98], esophagus [99], colorectal [100], ovarian [101] and lung [102] cancers have been documented. Association of upregulated genes in an altered group such as MCM10 [103], UBE2C [104–106] and CBX2 [107,108] with cancer progression corroborates impact of m^6^A pathway dysregulation on tumorigenesis. Significant upregulation of the cancer-related genes in the patients’ samples with genetic alteration in the m^6^A pathway provides another layer for evidence to support the link between the progression of BIC and the m^6^A pathway. This notion is well-preserved for the biological processes enriched by significantly upregulated genes in the altered group. As the most significant BPs such as ‘nuclear division’, ‘chromosome segregation’, ‘mitotic nuclear division’, were related to cell division, it is reasonable to suggest that genetic alterations in the m^6^A pathway may have an imposing effect on BIC progression. On the other hand, top significantly down-regulated genes in the altered group enriched the BPs containing ‘tubulin binding’, ‘microtubule binding’ and ‘ATPase activity’ important for the maintenance of cellular activities. Very similar findings obtained from the KEGG and GSEA (Figure S5) for the list of genes in altered and unaltered groups further confirm the over-representation enrichment analyses.

We next construct the PPI network between the most significant BPs and m^6^A pathway to reveal the linker proteins in these BPs directly interacting with m^6^A pathway components. Cell cycle-related proteins UBE2C, CCNE1, CCNE2, and PKMYT1 were identified as linker members of ‘nuclear division’ pathway with m^6^A regulatory proteins, and GLRB and MAPT proteins were ‘regulation of membrane potential’ to the m^6^A pathway (Figure 12). Initiation or progression of various cancers such as breast, gastric, ovarian, and lung cancers was accompanied by over-expression of G1/S-specific cyclin E1 and E2 [109–112]. Protein kinase membrane-associated tyrosine/threonine 1 (PKMYT1) can play tumor-suppressor or oncogenic roles and its expression in many cancers is elevated as well [113,114]. Hence, tumorigenicity-associated roles of connector proteins belonging to the nuclear division process are noticeable.

Strong association of genetic alterations in the m^6^A pathway and BIC progression based on the performed analysis cued us to extend our examination comprising the m^5^C and m^1^A pathways. A similar altered vs unaltered comparison regarding the genetic alterations in these pathways was carried out as done for the m^6^A pathway. Very remarkably, cell cycle-related processes such as ‘chromosome segregation’, ‘nuclear division’, ‘mitotic nuclear division’, and ‘sister chromatid segregation’ were detected as enriched BPs in the altered group of m^5^C or m^1^A pathways (Figure 8). These findings extrapolated the common basis of different epitranscriptomic pathways to progress the BIC by modulating the expression level of the genes taking essential roles in proliferation and cell cycle. Therefore, from our findings, we can speculate that aberrant expression of epitranscriptomic pathways results in upregulation of cell cycle and anti-apoptotic related genes and thereby increase the cell survival and proliferation rate of cancer cells.

Finally, to place the last piece of the puzzle, we investigated the common and unique BPs affected by the genetic alterations in m^6^A, m^5^C, or m^1^A pathways (Figure 9) to assess the commonality in downstream processes modulated by these pathways. Among the high number of shared BPs enriched by upregulated genes, surprisingly, most of them were found as cell cycle and proliferation-related processes which indicates the overlap of proto-oncogenes as upregulated DEGs in downstream processes. We observed that, within the BIC samples, genetic alterations of RNA modifying genes eventually lead to significant upregulation of survival and cell division-related genes and most probably account for progression and poor prognosis of BIC. Correspondingly, downregulated genes enriching BPs largely differed among m^6^A, m^5^C, or m^1^A pathways. Identified unique BPs for each epitranscriptomic pathway can be classified in the non-cancer-related pathways and this may underline the fact that besides the tumorigenic pathways, other cellular BPs are divergently affected by the genetic alterations in m^6^A, m^5^C, or m^1^A pathways.

Overall, the findings of this study support the link between dysregulation of RNA modifications and BIC initiation and progression. In accordance with previous studies, our data demonstrated that several elements of RNA modification pathways might be adopted as promising predictive biomarkers for BIC progression. Furthermore, the association of genetic alterations in epitranscriptome pathways with cell division and cell cycle-related BPs uncover the RNA modifying genes as candidate target genes for more effective treatment of BIC.

## 5. Conclusions

Our study has demonstrated the critical impact of genetic alterations in RNA modification pathways on BIC progression. Harnessing publicly available datasets and online databases, the presented study provides a comprehensive analysis of how the m^6^A pathway is associated with BIC pathogenesis and progression. We demonstrated that cell cycle, proliferation, and anti-apoptotic genes are particularly affected by the dysregulation of epitranscriptomic pathways. We anticipate that the generated data in this study is a useful catalog for the research community to design large-scale wet-lab experimental studies to confirm the therapeutic and prognostic significance of RNA modification genes in breast cancer treatment and prognosis.

## Supplementary Materials

The following are available online at www.mdpi.com/xxx/s1, Figure S1: Expression levels of m^6^A RNA modification regulatory genes in Breast Invasive Carcinoma pathological grades, Figure S2: Analysis of promoter methylation levels of m^6^A modification regulatory genes in Breast Invasive Carcinoma, Figure S3: Methylation status of m^6^A modification regulatory genes in Breast Invasive Carcinoma pathological grades, Figure S4: Kaplan-Meier survival curves of m^6^A regulatory genes in Breast Invasive Carcinoma regarding the mutation status, Figure S5: Genetic alterations in m^6^A pathway impact on KEGG pathways and gene ontologies based on GSEA.

## Author Contributions

Conceptualization, T.D.; methodology, T.D.; formal analysis, T.D., S.A.; investigation, T.D., S.A., M.Y.; writing—original draft preparation, T.D, M.Y.; writing—review and editing, T.D.; visualization, T.D., S.A., M.Y.; supervision, T.D; All authors have read and agreed to the published version of the manuscript.

## Institutional Review Board Statement

Not applicable.

## Informed Consent Statement

Not applicable.

## Data Availability Statement

All data analyzed and generated during this study retrieved from publicly available TCGA database (http://cancergenome.nih.gov) for Breast Invasive Carcinoma (TCGA, Pan Cancer Atlas).

